# Efficient Prospective Electric Field-Informed Localization of Motor Cortical Targets of Transcranial Magnetic Stimulation

**DOI:** 10.1101/2025.02.20.639076

**Authors:** David Luis Schultheiss, Zsolt Turi, Srilekha Marmavula, Peter Christoph Reinacher, Theo Demerath, Jakob Straehle, Joschka Boedecker, Matthias Mittner, Andreas Vlachos

**Affiliations:** Neurobotics Lab, Department of Computer Science, University of Freiburg, Freiburg, Germany; Department of Neuroanatomy, Institute of Anatomy and Cell Biology, Faculty of Medicine, University of Freiburg, Freiburg, Germany; Department of Stereotactic and Functional Neurosurgery, Medical Center - University of Freiburg, Faculty of Medicine, University of Freiburg, Freiburg, Germany; Fraunhofer Institute for Laser Technology (ILT), Aachen, Germany; Department of Neuroradiology, Medical Center-University of Freiburg, Faculty of Medicine, University of Freiburg, Freiburg, Germany; Department of Neurosurgery, Medical Center-University of Freiburg, Faculty of Medicine, University of Freiburg, Freiburg, Germany; Institute for Psychology, Norwegian University of Science and Technology, 7491 Trondheim, Norway; Institute for Psychology, UiT-The Arctic University of Norway, 9019 Tromsø, Norway; Center for Basics in NeuroModulation (NeuroModulBasics), Faculty of Medicine, University of Freiburg, Freiburg, Germany; Center BrainLinks-BrainTools, University of Freiburg, Freiburg, Germany

**Keywords:** Transcranial magnetic stimulation, electric field, motor mapping, motor evoked potentials, motor cortex, farthest point sampling

## Abstract

Transcranial magnetic stimulation (TMS) is a versatile non-invasive tool for brain mapping and neuromodulation in both healthy individuals and patients. Effective TMS-based causal brain mapping relies on precise localization of cortical targets. Current state-of-the-art approaches use statistical methods to quantify the relationship between TMS-induced electric fields (E-fields) and motor evoked potential (MEP) amplitudes. However, this method typically relies on the random selection of coil configurations, which limits its efficacy. In this study, we present a novel optimization strategy for TMS-based motor mapping by prospectively selecting coil configurations based on their E-field characteristics using an iterative sampling algorithm called farthest point sampling (FPS). Through a combination of theoretical analysis, simulation and experimental validation including 10 healthy individuals, we systematically evaluated the performance of FPS against the random sampling approach. Our results demonstrate that FPS is twice as efficient as random sampling in reducing the number of trials required for estimating the motor map, while also being more robust across participants and less susceptible to noise. These findings highlight the potential of FPS to significantly enhance the efficiency of motor mapping, paving the way for the development of more effective TMS mapping algorithms.

## 1 Introduction

Transcranial magnetic stimulation (TMS) is a widely used interventional technique for the non-invasive stimulation of neural tissue in humans and animals [1; 2; 3; 4]. TMS generates short-lasting intracranial electric fields (E-field) of approximately 200-300 *µ*s duration (for bipolar pulses), reaching a strength of up to 150-200 V m^−1^ in the human neocortex [1; 5; 6; 7; 8]. This E-field can depolarize neurons and, under certain conditions, trigger action potentials in specific subset of neurons [9]. TMS has emerged as a valuable tool for brain mapping in both neuroscience research and clinical applications, such as pre-surgical planning [10; 11; 12; 13; 14; 15; 16].

Brain mapping aims to establish causal links between brain regions and their associated functions. However, this task is complicated by the hierarchical organization of the brain, where brain regions are organized within larger functional networks [17; 18; 19]. Effective brain mapping requires the precise localization of cortical target–specific areas where TMS elicits measurable neuronal effects causally linked to an induced biological response. For instance, TMS applied to the human primary motor cortex can evoke muscle twitches in contralateral peripheral muscles, measurable as motor evoked potentials (MEPs) [1; 10; 20]. Despite the widespread use of TMS-induced MEPs, the precise regions in the motor cortex responsible for these effects remain a topic of ongoing investigation [9; 21; 22; 23; 24; 25; 26; 27], limiting the effectiveness of TMS as a brain mapping tool. An improved understanding of how TMS impacts cortical targets can potentially enhance both neuroscience research and its application in clinical situations.

Various methods have been developed for TMS-based functional mapping of the motor cortex. These methods often focus on delineating motor cortical representations of peripheral muscles (hereafter referred to as muscle representations), such as the first dorsal interosseus muscle (FDI) [10; 28]. The conventional grid method involves stimulating a predefined grid of scalp sites, followed by calculating the center of gravity based on the evoked motor responses [10; 29; 30; 31]. However, this method estimates muscle representations using locations correspondig to the scalp and cannot accurately capture their spatial dispersion within the motor cortex with the desired level of precision (cf. [26]).

State-of-the-art approaches aim to improve localization by statistically quantifying the relationship between the TMS-induced E-field and MEP amplitudes [26; 28; 32]. These methods typically involve the collection of MEPs generated by hundreds of stimulations at randomly selected coil positions, followed by retrospective E-field simulations at each location [26; 28; 32]. These approaches feature significant improvements both in terms of efficiency and spatial resolution compared to grid-based methods. However, since the random selection of stimulation site and coil location does not prioritize the most informative coil configuration, it can potentially take a large number of trials before the method converges on a solution with satisfactory precision. In particular, randomly selected coil positions frequently generate redundant E-field characteristics, unnecessarily increasing the number of trials needed for the mapping. Therefore, identifying optimal coil configurations needed for efficient and precise motor mapping is critical, particularly in clinical settings, where minimizing the experimental burden on patients is essential.

This study presents a novel, prospective E-field-informed TMS-based motor cortical mapping method, designed to improve efficiency. By employing the sampling algorithm *farthest point sampling* (FPS), we iteratively select maximally informative coil configurations based on their E-field characteristics. Our theoretical and experimental analyses congruently demonstrate that FPS significantly reduces the number of trials required for motor mapping compared to random sampling, offering a more efficient approach beneficial for both research and clinical applications.

## 2 Methods

### 2.1 Study overview

In this study, we systematically evaluate the performance of the random sampling and the FPS algorithm through both, theoretical and experimental work. In the theoretical section, we develop a synthetic MEP generation model for in-silico simulations, allowing us to evaluate the performance of the two algorithms across various scenarios. In the experimental part, we test the performance of both algorithms on healthy individuals using real-time neuronavigated robotic-arm guided TMS.

### 2.2 FPS algorithm

#### Algorithm 1

Farthest Point Sampling

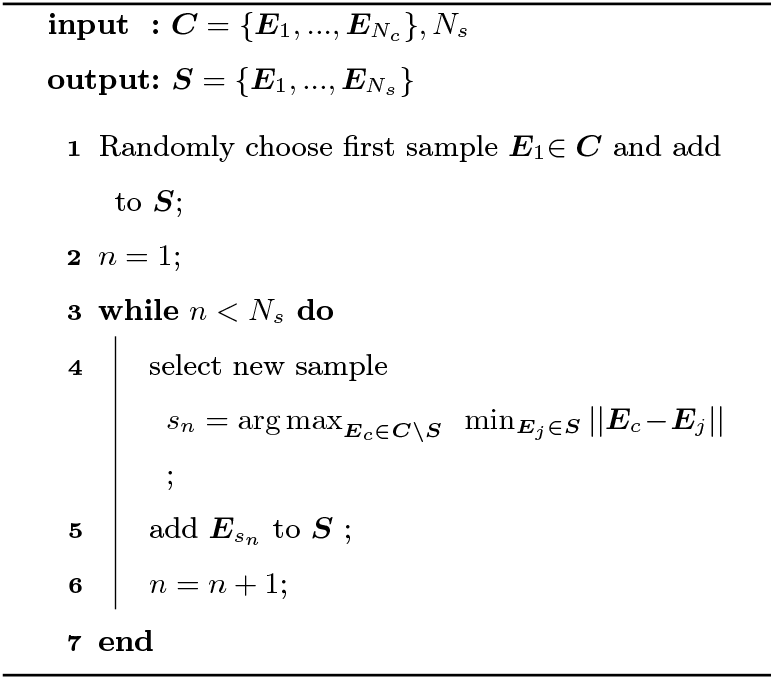

The goal of the experiment is to prospectively select coil configurations informed by their induced E-field properties within the region of interest (ROI). For a given coil configuration *c*, the simulated ROI E-field map is represented as ***E***_***c***_. The configuration for each pulse is a unique combination of coil location *x*_*c*_, *y*_*c*_ and coil orientation *α*_*c*_. Denoted as ***θ***_*c*_ = *{x*_*c*_, *y*_*c*_, *α*_*c*_*}*. Since the motor mapping experiment is limited to approximately a few hundred TMS trials per experimental session, a sampling algorithm is required to select a small subset 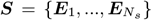 from a large candidate dataset 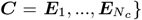.In theory, the size of the candidate set is infinite as coil location and orientation span a continuous parameter space. However, similar configurations produce similar E-field distributions and we hypothesize that in order to decorrelate candidate locations in the ROI, dissimilar E-field distributions are required. Thus, the parameter space is discretized, which allows sampling from a finite set of *N*_*c*_ E-field candidates. Depending on the resolution of the potential coil center distances and orientation angles, this could result in millions of coil configurations and, consequently, millions of potential E-field distributions. However, as shown in Appendix A, coarser resolutions suffice, reducing *N*_*c*_ to 1017.

A well-known and data-efficient iterative sampling technique is FPS [33; 34]. The idea of FPS is to select a small subset from a larger dataset while ensuring that the selected samples are representative of the original set. In the present case, we choose sequential coil configurations that generate maximally different E-fields in the ROI.

The first sample is chosen randomly from ***C***. Then, the algorithm iteratively adds E-fields to the subset by choosing the E-field that is maximally dissimilar from the ones already selected. Hereby, 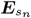 is considered to be maximally dissimilar at step *n*, if its minimal Euclidean distance to previously sampled E-fields is maximized across all remaining candidate E-fields. This can be formulated as the following maximization condition for selecting *s*_*n*_:

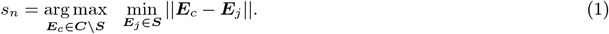

Here, ||.|| denotes the Euclidean norm. A pseudocode of FPS is presented in Algorithm 1.

### 2.3 Synthetic MEP simulations

We conducted a comprehensive simulation-based analysis based on synthetic MEPs to evaluate the performance of the two algorithms. Synthetic MEP simulations were based on a biologically inspired MEP generation map (MGM) and an MEP model. This simulation setup allows us to i) sample MEPs for arbitrary coil configurations, ii) place hypothetical muscle representations at different anatomical locations in motor cortex (e.g. gyral crown, gyral lip, sulcal wall, center of ROI, edge of ROI), and iii) study the effect of noise on the estimation of the motor map.

#### MEP generation map

We hypothesize the existence of a spatially restricted muscle representation in the motor cortex which is responsible for generating MEPs in that specific muscle. In our high-resolution representation of the human head, the muscle representation is discretized into *K* discrete compartments. These compartments are triangles if the ROI is defined on the brain surface or tetrahedrons if the ROI is a brain volume. For our simulations, we position the muscle representation at any compartment *k*_max_ ∈ *K*, referred to as the most excitable compartment. Since the exact size and boundaries of the muscle representation in human motor cortex are uncertain, we model excitability as a distribution centered around the most excitable location *k*_max_. A Gaussian probability density function, which decays exponentially with distance from its center, is a suitable choice for this distribution. Based on this, the excitability *m*_*k*_ of compartment *k* is defined as

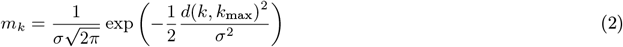

where *d*(*k, k*_max_) is the geodesic distance on the cortex of compartment *k* from the most excitable location *k*_max_. The spatial extend of the excitable area is controlled by the standard deviation of the Gaussian distribution *σ*. The excitabilities of all compartments are summarized in the MGM ***M***. This approach assumes that excitability decreases smoothly as the distance from the most excitable location increases, forming a focused excitability profile.

#### MEP model

An underlying assumption guiding motor mapping within the primary motor cortex is the presence of a sigmoidal relationship between stimulation strength *x* in an excitable location of the cortex and MEP amplitude *y* [28; 32]. The sigmoid function takes the form

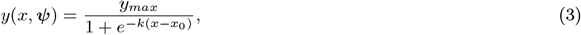

where *x*_0_ is the location of the turning point on the abscissa, *y*_*max*_ is the saturation amplitude, and *k* is the slope. The parameters are summarized in ***ψ*** = *{x*_0_, *y*_*max*_, *k}*.

We define the stimulation strength *x* as the inner product of the MGM ***M*** and the stimulus E-field ***E***:

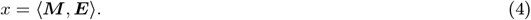

This reflects the assumption, that strong E-fields that overlap the muscle representation will increase the measured MEP while activations outside the MGM will not affect it. Then, zero-mean Gaussian noise *ϵ* is added to mimic variability stemming from a variety of sources (e.g., small coil or head movements, brain-state, etc.) inherent to experimental measurements, ensuring that the simulated data more accurately reflect real-world conditions:

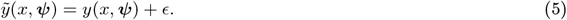

We observed in previously recorded data that noise tends to be stronger for a higher stimulation intensities. However, it also seems to saturate for very strong E-fields. Therefore, the noise is modulated by a sigmoid term and noise amplitude *P* and *ϵ* → *ϵ*(*x*, ***ψ***, *P*) with

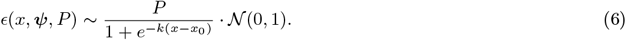

This ensures that noise saturates to *P* as the stimulation strength becomes maximal and that the noise amplitude goes to zero as *x* gets smaller. Additionally, the noisy MEPs are clamped to have minimum 0, as peak-to-peak MEPs can only be positive. The MEP generation process is illustrated in Figure 1.

**Figure 1:**
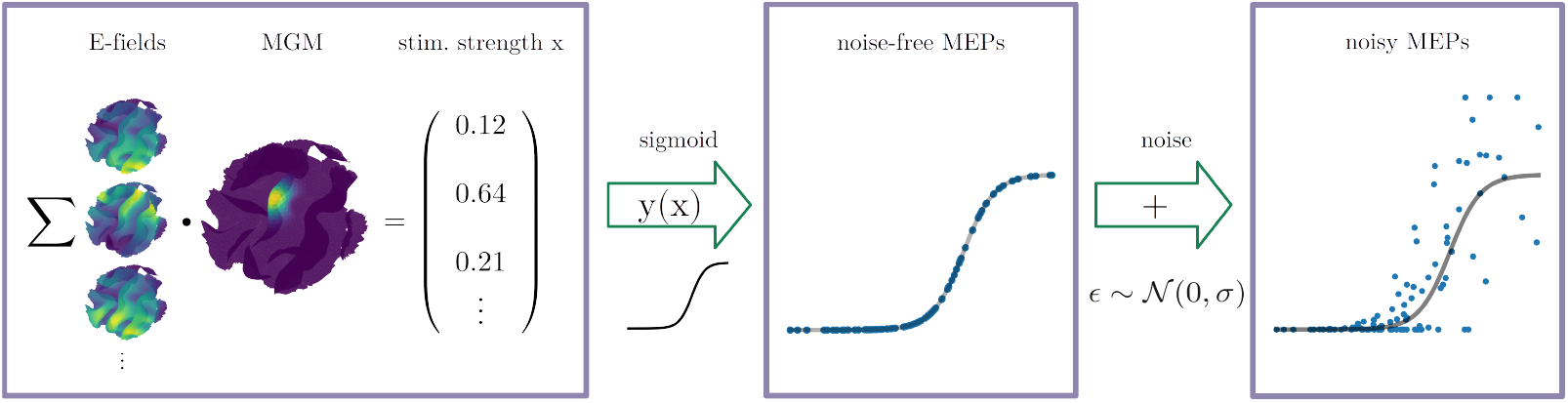
Synthetic MEP simulations. The figure depicts the process of generating synthetic MEP amplitudes. The inner product of a user defined MGM and E-fields (one per coil configuration) result in the latent stimulation strength *x*. Then, *x* is passed through a sigmoid function. Lastly, Gaussian noise is added to retrieve the variability inherent in experimental measurements of MEP amplitudes. The MEP amplitudes are clipped to 0 as lower bound.

### 2.4 Motor mapping of synthetic and experimental data

For generating muscle representation maps in the motor cortex, we closely follow [28; 32]. At step *s* of the sampling procedure, there are *s* E-field values per ROI compartment in the sampled dataset. For each sample, we also have a corresponding MEP amplitude, that was either experimentally measured, or synthetically generated by the model from eq. 5. Then, for each compartment *k*, a sigmoid function is fitted to the MEP data using the Levenberg-Marquardt least-squared optimizer [35]. The resulting *R*^2^ values for all compartments are then combined to construct an *R*^2^-map ***R***^***2***^ for the ROI [32].

The fitting score score_*f*_ is then computed as the inner product of a reference *R*^2^-map 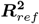,e.g. ***M*** or a high resolution map fitted with all samples in ***C***, and an estimated *R*^2^-map 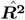,obtained from the small subset of samples selected using random or FPS sampling. Both maps are normalized to have *l*^2^-norm 1:

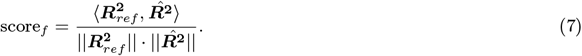

The score can be interpreted as the similarity or overlap of the two maps where a score of 1 is a perfect fit and 0 is the lower bound.

### 2.5 TMS experiment

#### Ethics declarations

The Ethic Committee of the University of Freiburg Medical Center approved the investigation (application number: 23-1114-S1). All participants gave written informed consent before participation. We performed all experiments in accordance with relevant guidelines and regulations.

#### Participants

Ten healthy, right-handed participants (6 females, 4 males; age [years]: mean*±*std: 28.8 *±* 1.62, range from 21 to 26) were recruited for the study. Participants had laterality index of mean*±*std: 83.49 *±* 16.45, range from 57.14 to 100, as determined by the Edinburgh Handedness Inventory [36]. The selection process for volunteers began with a personal meeting with one of the study physicians, during which the exclusion and inclusion criteria were reviewed and approved. Next, participants who expressed interest in the study and met all inclusion criteria, while also not meeting any exclusion criteria as evaluated by the study neurologists, underwent structural MRI acquisition. The individual MRI images were then examined by the study radiologist to identify any brain pathologies. No participants showed signs of brain pathology and all of them were therefore eligible to be included in the study. In addition, we applied further exclusion criteria for data included in the analyses. One participant was excluded due to an incorrect search radius of 50 mm instead of the intended 30 mm (see section *Head modeling and macroscopic E-field simulations* for details). Another participant was excluded due to delayed MEPs occurring outside the predefined time window of 18 to 35 ms following TMS application [26]. Therefore, of the 10 volunteers, data from eight were used in the motor mapping analysis.

#### MRI

High resolution MRI data was acquired with a 3-Tesla SIEMENS MAGNETOM Prisma scanner and a 64channel head coil. T1-weighted images were acquired with the following parameters: sequence: Magnetization Prepared - RApid Gradient Echo (MPRAGE); voxel size: 1.0 *×* 1.0 *×* 1.0 mm; TR: 2500 ms; TE: 2.82 ms; flip angle: 7 ^*°*^; FoV: 256 mm; fat suppression: water excitation; orientation: sagittal; no. of slices: 192; total acquisition time: 3 min. 58 s. T2-weighted images were acquired with the following parameters: sequence: Sampling Perfection with Application optimized Contrasts using different flip-angle Evolutions (SPACE); voxel size: 1.0 *×* 1.0 *×* 1.0 mm; TR: 2500 ms; TE: 231 ms; FoV: 256 mm; fat suppression: no; orientation: transversal; no. of slices: 160; total acquisition time: 6 min. 42 s. In addition to the high-resolution 3D anatomical images, we also acquired diffusion tensor imaging data with the following parameters: voxel size: 1.5 *×* 1.5 *×* 3.0 mm; TR: 2800 ms; TE: 88 ms; no. of directions: 65; FoV: 222 mm; total acquisition time: 6 min. 22 s.

#### Head modeling and macroscopic E-field simulations

We used an open access toolbox called Simulation of Non-invasive Brain Stimulation (SimNIBS; version 3.2.6) to create head models and perform E-field simulations [5]. Anatomically realistic multi-compartment head models were created by calling the *headreco* pipeline using individual T1- and T2-weighted anatomical and diffusion-weighted MRI sequences. To enhance the smoothness of the skin compartment, we individually tailored the number of cortical smoothing repetitions, ranging between 100 and 200 repetitions [8]. The final head mesh contained around 4 million tetrahedra. For the following tissue compartments, we used the default isotropic conductivity values [in S/m]: eyes (0.5), scalp (0.465), bone (0.01), cerebrospinal fluid (1.654), whereas for the gray matter and white matter compartments, we assigned volume normalized anisotropic conductivities [37]. E-field calculations for each stimulation were conducted with high resolution anisotropic finite element models. For all simulations, we set the stimulation intensity at a coil-current rate of change of 1.49 A/µs that corresponded to 1% MSO for our TMS device. The scalp-to-coil distance was determined individually by measuring hair thickness using a depth gauge [38; 39]. Additionally, an extra 1 mm was added to the scalp-to-coil distance to account for the touch sensor of the TMS coil. We established the ROI center based on a prior meta-analysis [40]. Specifically, we converted the FDI coordinates from the Montreal Neurological Institute coordinate space (MNI152; x-37, y-21, z+58) into the subject-space. The primary coordinate was subject to individual visual inspection, and where necessary, adjustments were made to ensure its localization within the precentral gyrus. Then, we calculated the Euclidean distance between the individual FDI coordinate within the gray matter compartment and the scalp compartment, choosing the scalp position with the closest proximity as the stimulation target. In the E-field simulations, we employed a search radius of 30 mm around the ROI center, positioning coil locations with a spatial resolution of 3 mm. For each cortical target, we systematically adjusted the coil’s rotation angle in increments of 5° from 0° to 175°. We excluded orientations at and beyond 180° since they predominantly affected the E-field direction. The constraints on these parameters are based on extensive preliminary simulations, as described in *Appendix A*. To obtain E-field values, we extracted element-wise data from the middle gray matter compartment using a cylindrical ROI with a radius of 20 mm from the ROI center [32]. Therefore, we utilized two radii around the ROI center. One radius, set at 30 mm, defined the coil search radius, while the other radius corresponded to the actual ROI used for analyzing the E-field. A larger radius was applied for the coil search to further enhance differences in the E-field distributions within the ROI, particularly between the center and edge regions.

#### Neuronavigated robotic-arm-guided TMS

We employed a neuronavigated, robotic-arm-guided TMS system in the experimental phase. Single biphasic TMS pulses were administered through a MagPro X100 stimulator (MagVenture, firmware version: 7.2.4) using a robot-compatible Cool-B65 CO figure-of-eight-coil. The Localite neuronavigation system (TMS Navigator Robotic Edition Axilum cobot; version 3.3.8) and an Axilum Cobot were employed for precise coil guidance (version 1.3.2.0; controller version 2.1.10.148). Matsimnibs files, corresponding to individual E-field simulations and containing the 4-D affine transformation matrices that define coil positions and directions, were converted to HTML files compatible with the neuronavigation system. These files were then uploaded to the neuronavigation software using Localite’s Session Utility Tool (version 3.3.15). The coil position selection was controlled by the data collector by using Localite’s neuronavigation software.

MEPs were recorded from the participants’ right FDI muscle. We placed one self-adhesive surface electrode over the FDI muscle belly and another one at the proximal interphalangeal joint (Kendall, H124SG, Cardinal Health Germany 507 GmbH, Norderstedt, Germany). The surface electrodes were connected to the amplifier system (D-360, Digitimer Ltd., UK, Welwyn Garden City) with bandpass filtering ranging from 10 Hz to 2 kHz. The amplifier system was subsequently connected to a Hum Bug Noise Eliminator (Digitimer Ltd., UK, Welwyn Garden City), which was then linked to a data acquisition interface (Power1401 MK-II, CED Ltd., Cambridge, UK) with a sampling rate of 5 kHz. Participants wore earplugs with a single-number rating isolation value of 35 dB during the experiment to attenuate the click sound generated by TMS (Moldex Spark Plugs 7800, Walddorfhäslach, Germany).

The synchronization between the TMS device and EMG data acquisition system was accomplished through a custom MATLAB application developed in-house. For controlling the time and the intensity of stimulation, we utilized the MAGIC toolbox [41]. Raw EMG data from the Power1401 MK-II data acquisition unit was directly accessed using the “Matced” interface (available at https://ced.co.uk/downloads/contributed). All incoming data was stored by the MATLAB application irrespective of whether a trial was later included or excluded from analysis. To ensure the integrity and reliability of the study, trials were not subject to deletion during online data collection. Trials were automatically excluded (but not deleted) from the data analysis if they displayed pre-stimulus muscle activity with a peak-to-peak amplitude exceeding 50 *µ*V within a 100 ms time-window before the application of TMS. The peak-to-peak amplitude of MEPs were automatically extracted from 18 to 35 ms time window after the TMS pulse [28]. In the resting motor threshold (RMT) estimation, we used a minimum peak-to-peak amplitude of 50 *µ*V. Together with the neuronavigated TMS-cobot, our MATLAB application reduces reliance on operator discretion for subjective decisions related to free-hand coil positioning, stimulation intensity determination, and online trial exclusions under limited time. Consequently, our systematic approach ensures an accurate and reproducible experimental workflow, prioritizing both participant safety and research integrity.

RMT was determined utilizing a maximum likelihood estimation based adaptive threshold hunting algorithm, following the recommendations of the International Federation of Clinical Neurophysiology [42]. This algorithm employs parameter estimation through sequential testing, commonly referred to as the PEST method [43]. Initially described by Awiszus [44], it was subsequently implemented as desktop software known as the Motor Threshold Assessment Tool (MTAT; version 2) [45] (but see references [46] and [47] for a recent update of MTAT to version 2.1). The current study utilized the MATLAB version of MTAT, obtained through personal communication with Julkunen [48]. The algorithm yields an accurate estimation of the RMT within 13 to 30 trials. The convergence of the RMT estimate, along with the estimated threshold and the corresponding MEPs, was verified for each estimation procedure by the data collector.

During motor mapping, each participant was planned to receive a total of 300 stimulations, consisting of 150 coil configurations selected using random sampling and 150 using the FPS method. However, the actual number of stimulations was slightly higher due to the need to repeat trials caused by muscle preactivity. The random and FPS stimulations were further divided into blocks of 50 TMS trials each. The order of random and FPS blocks was counterbalanced across participants: five participants started with the random blocks and finished with the FPS blocks, and vice versa. Before assigning the FPS trials to the three TMS blocks of 50 TMS trials each, the order of the FPS trials were randomized. This step was not necessary for the TMS trials generated by the random method. Motor mapping was performed at 120% of the RMT. However, for two participants, a reduced intensity of 110% RMT was used to minimize stimulation-induced subjective discomfort.

### 2.6 Research integrity

The code for conducting the E-field modeling, simulations and analyses in the present study is available for download at the following Github repository https://github.com/david-schu/efield-informed-motor-mapping. Data generated during the simulations were not archived due to their substantial size.

## 3 Results

### 3.1 Motor Mapping on synthetic data

First, we tested how well a ground truth motor map could be estimated depending on the location of the muscle representation. To do this, we placed *k*_*max*_ of the MGM ***M*** at every possible compartment *k* of the ROI, generated MEP amplitudes for a dense grid of 2,853 coil configurations, and computed 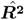 based on the respective E-fields and MEP amplitudes. Next, we calculated the overlap score, score_*f*_ 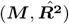,for each compartment. Figure 2 shows that there is no clear trend when comparing locations on the gyral crown or lip to those on the sulcal wall. Only locations within the gyral fold exhibit consistently low fitting scores across the entire ROI.

**Figure 2:**
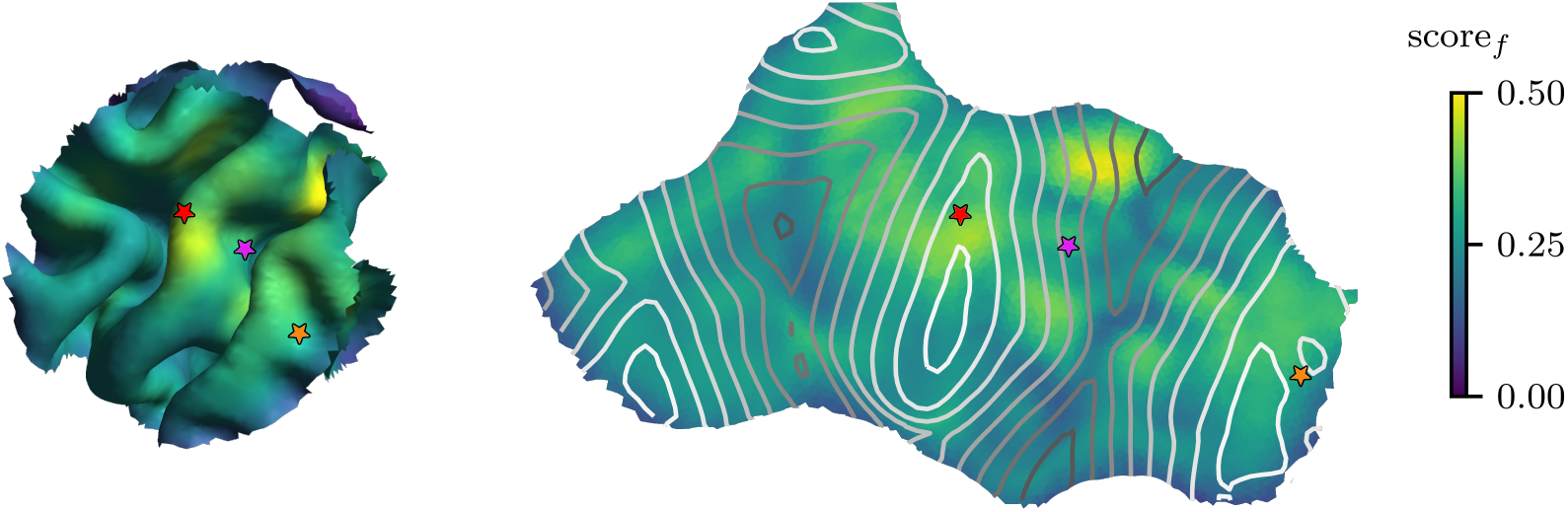
Optimal fitting scores depend on location of muscle representation. We placed the muscle representation at every possible element in the ROI and tested how well the ground truth motor map could be recovered with a fitted *R*^2^-map, computing score_*f*_ 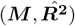 for each location. (A) The scores mapped to a three-dimensional representation of the ROI. (B) The three dimensional ROI surface was flattened with multidimensional scaling to obtain a two dimensional representation based on the pairwise matrix of geodesic distances between ROI compartments. The contour lines indicate sulcal depth, with light colors corresponding to the gyral crown and darker colors to the sulcal fold. The colored stars mark identical locations in both ROI representations.

To evaluate the efficacy of sampling methods for motor mapping, we compared their ability to fit motor maps to synthetic MEP amplitudes generated from predefined motor maps ***M***. The MEP amplitudes were generated as described in Section 2.3. The experiment was run on a single head model and repeated using 10 random sampling seeds and 10 random seeds for generating MEP noise, resulting in a total of 100 runs per parameter setting.

Figure 3 A1-C1 show that the ground truth MGM ***M*** is recovered much faster with FPS compared to random sampling. While both methods converge to similar scores for large sample sizes, FPS is more data efficient in the early sampling stages. Overall, the scores indicate that muscle representations located at the ROI center identified with higher precision compared to those at the ROI edge. The same can be observed for the target being at gyral crown compared to the sulcal wall or fold. However, even at the ROI edge, target localization was feasible, particularly with FPS-guided stimulation, underscoring the robustness of FPS for motor mapping.

**Figure 3:**
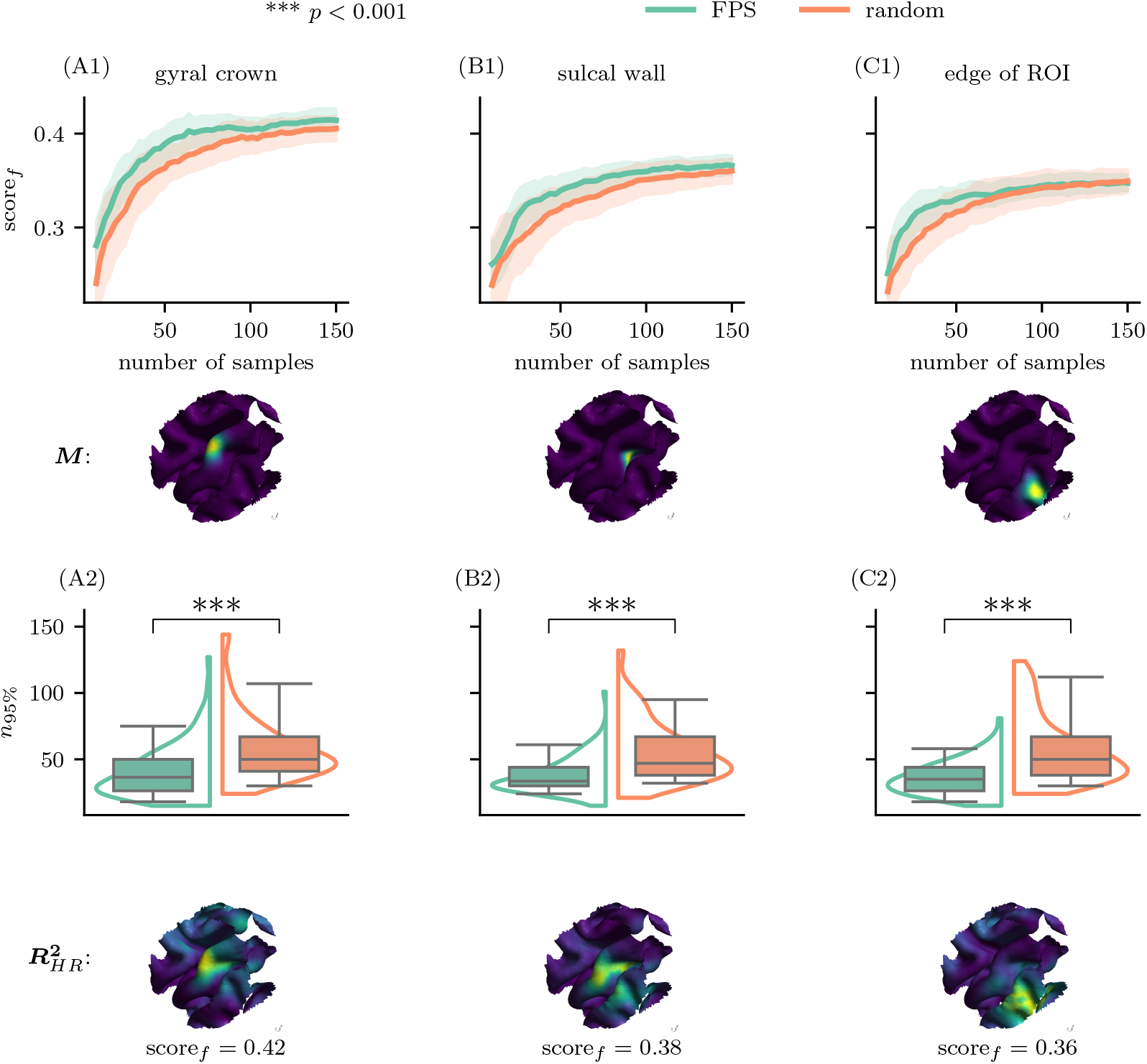
Subsampling from the synthetic data demonstrates superior performance of the FPS algorithm compared to the random sampling across all simulation conditions. We subsampled from a large dataset of simulated E-fields. The motor maps ***M*** used to generate MEPs were placed at the gyral crown in the center of the ROI (A), the sulcal wall in the center of the ROI (B), and the gyral crown at the ROI edge (C). We tested how well ***M*** could be recovered by computing score_*f*_ with increasing number of samples (A1-C1). The lines represent the mean across 100 runs, the shaded areas the standard deviation from the mean. Next, we tested how many samples were needed to reach 95% overlap (score_*f*_ *>* 0.95) with 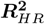 (A2-C2). The boxes represent 100 random initializations. The lines in the boxes represent the median, the whiskers extend from the 5th to the 95th percentile. The violins in the background show a fitted kernel density estimate of the data distribution. The asterisks above the plots represent significant differences between the number of samples needed to reach the target score with FPS and random sampling using a permutation test. The score below the illustration of each 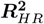 map is the overlap of 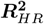 and the respective ground truth ***M***. FPS outperforms random sampling for all three simulated muscle representations.

To establish an upper performance limit, we created a high resolution (HR) reference map 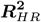 with ≈ 13000 MEP samples. This HR map represents the theoretical outcome if stimulation were applied to every point on a dense grid (spatial resolution 2mm, angular resolution 10^*°*^, search radius of 30 mm) and served as the ground truth target. The performance of each sampling method was quantified by determining the number of samples required to achieve a fitting score score_*f*_ of 0.95, corresponding to 95% overlap with 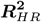. Figure 3 A2-C2 confirms that the 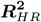 –maps were recovered more quickly and accurately using FPS compared to random sampling across all three tested simulated muscle representations. When the target was located at the center of the ROI on the gyral crown, FPS achieves superior performance, requiring an average of 40.39 *±* 19.96 (mean*±*std) samples, compared to random sampling 60.64 *±* 25.00. This results in a difference of ≈ 20 samples (Figure 3A). For a target at the center of the ROI but on the sulcal wall, the difference between the methods narrowed, with FPS requiring 37.42 *±* 13.30 samples compared to 55.20 *±* 22.50 for the random sampling, a reduction of ≈ 18 samples (Figure 3B). At the edge of the ROI, FPS demonstrates the largest improvement, requiring 35.98 *±* 12.87 compared to 59.15 *±* 24.61 for the random sampling, a reduction of almost 24 samples (Figure 3C).

On average, FPS required approximately 40 samples to achieve a fitting score of 95% with the 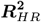 -map across all target locations. These results underscore the significant sample efficiency of FPS for motor mapping. Furthermore, FPS demonstrated reduced sensitivity to randomization of initialization, as indicated by smaller whisker extensions and lower standard deviations in Figure 3.

To assess the statistical significance of differences between the FPS and random sampling distributions, a permutation test was conducted. Using 10,000 permutations and the difference in means as the test statistic, the results yielded *p <* 0.001 for all target locations (Figure 3 A2-C2). Importantly, FPS consistently outperformed random sampling regardless of MEP model parameters ***ψ***. A comprehensive parameter sweep over MGM standard deviation *σ*, MEP generation noise *P*, and sigmoid slope *k* yielded comparable performance differences. For larger noise amplitudes *P* the scores generally dropped. The same was observed for smaller MGM standard deviations, equivalent to more focalized muscle representations. The change of the sigmoid slope *k* had no substantial effect on the results. These observations are described in more detail and also visualized in *Appendix B*.

### 3.2 Motor mapping on human participants

Building on the insights gained from synthetic data, we next evaluated the performance of the proposed sampling methods in motor mapping experiments conducted with human participants to assess their practical applicability and robustness.

No serious adverse effects (e.g., epileptic seizure, syncope, etc.) occurred during or after the experiment. Among the 10 participants, none reported perceiving phosphenes during the experiment. One participant experienced a mild headache during the experimental session, rated at the lowest intensity (1/10) on a Likert scale. Overall, participants described the stimulation pulses as a moderate tactile sensation on the scalp, with a mean score of 4.75 *±* 2.15 on a 10-point Likert scale (1 = minimal sensation, 10 = maximal sensation). For one participant with a mapping intensity at 79% MSO, the stimulation was perceived as uncomfortably intense at specific frontal and temporal locations. Participants also described the stimulation pulses as moderate auditory sensations with a mean score of 4.55 *±* 2.54 on a 10-point Likert scale.

For each participant, we generated a target *R*^2^-map, by fitting all measurements, as well as *R*^2^-maps based on data from the random and FPS block. These maps are shown in Figure 4. Subsets of the two blocks were then analyzed to determine how many stimulations were required to approximate the target map. Two metrics were evaluated: (1) the fitting score score_*f*_ between the target map and subset maps, as defined in Equation 7, which the similarity muscle representation distributions across the ROI surface; and (2) the geodesic distance *d* between the peak *R*^2^ value of the target map and the subset map, representing the distance between the muscle representation centers.

**Figure 4:**
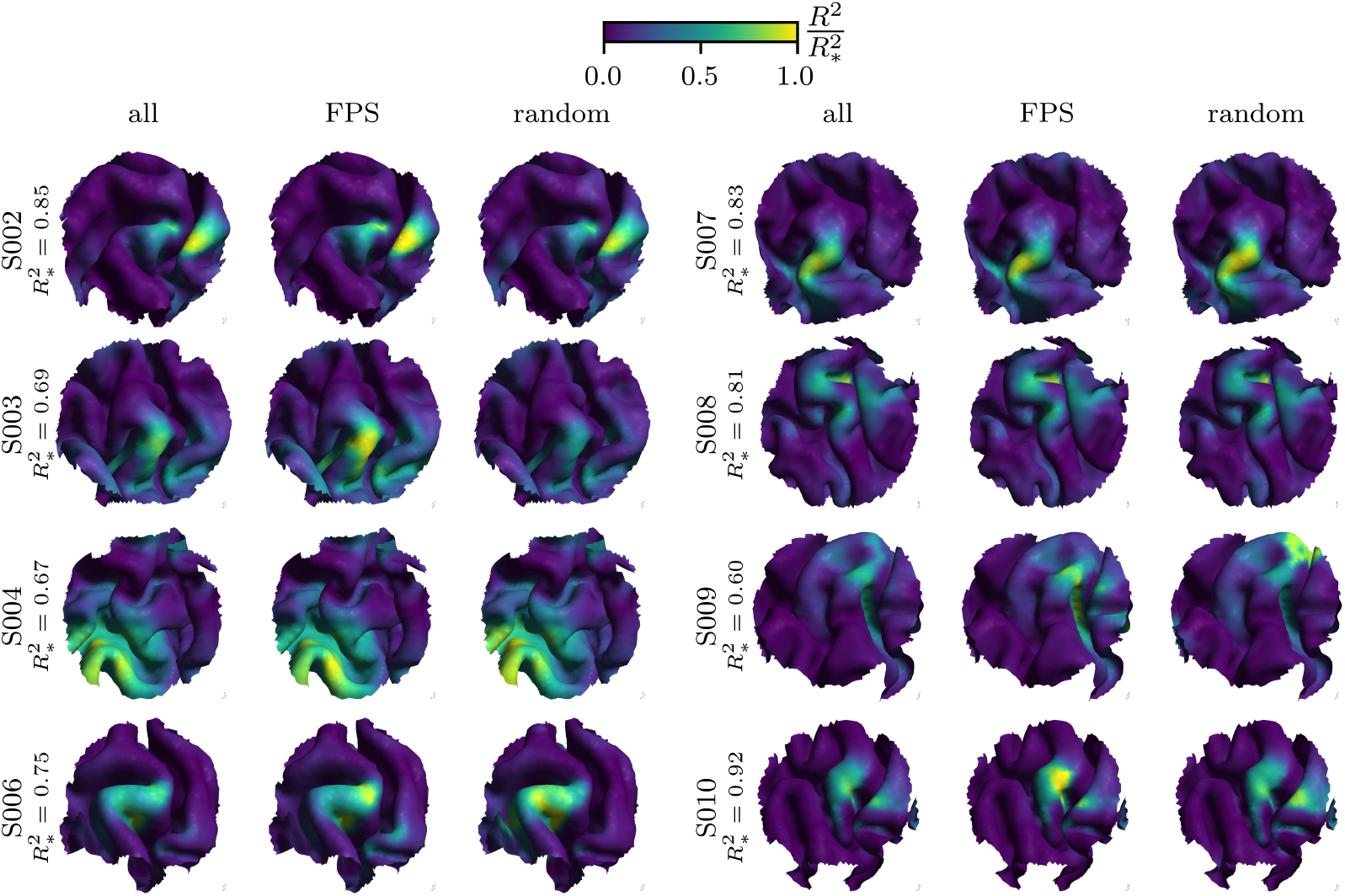
Predicted FDI motor cortical representations of all participants. For each participant that entered the analysis, the results given all samples, FPS samples, and random samples are depicted in the three columns. The *R*^2^-values are normalized by the subject-specific maximum value 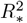.Theses maximal values range from 0.60 to 0.92. The *R*^2^-maps can be interpreted as the subject-specific muscle representation of the FDI muscle. The representation for the FDI muscle of most participants lies at the gyral crown of the precentral gyrus.

Figure 5 illustrates several key differences between FPS-based stimulation maps and random sampling methods stimulation maps. FPS-derived maps converged to the target map notably faster, with this advantage being most pronounced during the early stages of sampling (Figure 5A). The geodesic distance *d* between the highest *R*^2^ value of each sampled map and the target map was consistently smaller for FPS compared to random sampling, particularly at lower sample sizes (Figure 5B). Moreover, both score_*f*_ and the distance *d* exhibit greater robustness across participants in FPS-derived maps, as reflected by narrower 95% confidence intervals for these metrics. Notably, FPS achieved 95% overlap with the target map using ≈ 50 E-field MEP pairs, whereas random sampling required nearly twice as many samples to reach the same level of accuracy. To ensure generalizability of our findings, we conceptually replicated the analysis using an independent dataset previously published by et al. [32], as described in *Appendix C*.

**Figure 5:**
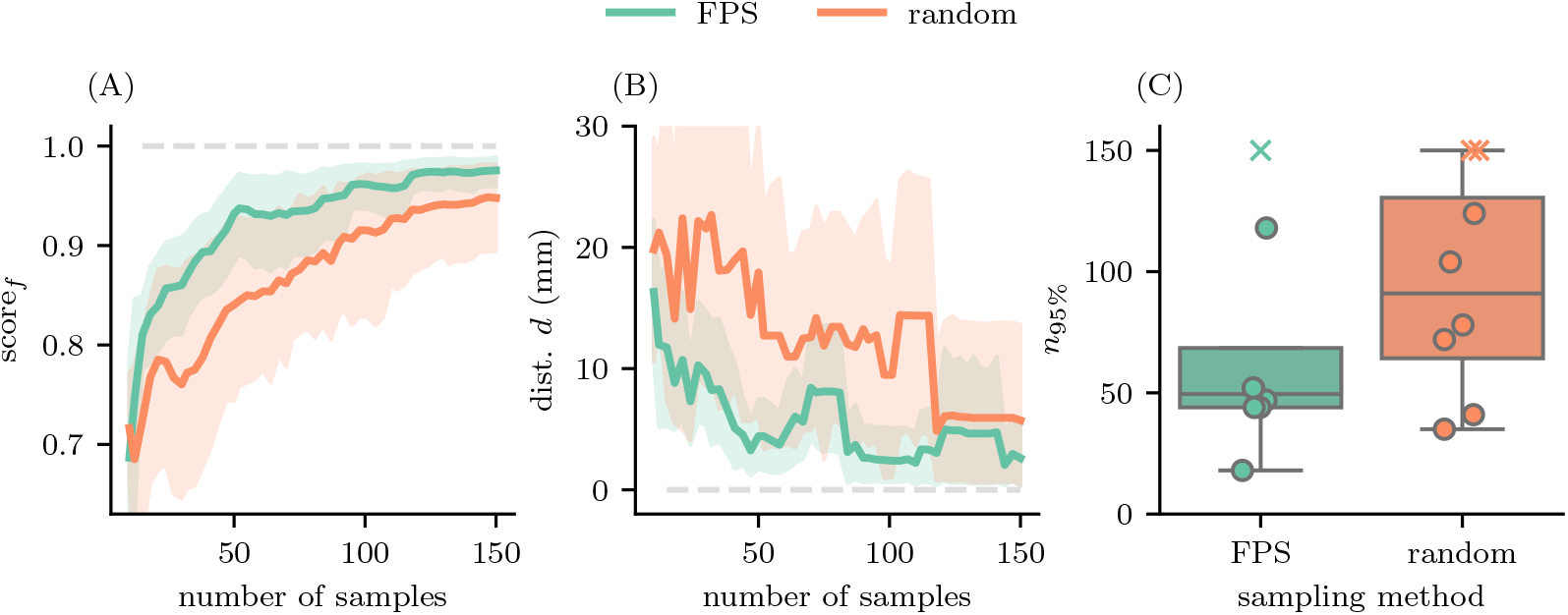
Subsampling from the experimental dataset. For every other sample we calculated (A) the fitting score score_*f*_ as overlap of the fitted motor maps with the *R*-map fitted with all valid samples (ideally 300), and (B) the geodesic distance *d* of the respective mesh elements with the highest *R*^2^-values. The lines represent the mean across participants and the shaded area the 95% confidence interval. (C) We tested how many samples were needed to reach 95% overlap with the target *R*^2^-map (score_*f*_ *>* 0.95). The boxes summarize *n*_95%_ across participants and extend from the 25th percentile to the 75th percentile. The line in the boxes represents the median, the whiskers extend from the 5th to the 95th percentile, the points represent single participants, and participants where score_*f*_ did not reach 95% are marked as crosses and placed at *n*_95%_ = 150. FPS outperforms random sampling especially in the early sampling stage and is approaching an optimal overlap and small target distance much faster.

## 4 Discussion

The current state-of-the-art approach to TMS-based motor mapping relies on the random selection of coil configurations during MEP assessments, followed by retrospective modeling of the induced E-field [26; 28; 32]. While this method has advanced the field significantly, it remains inefficient, especially in contexts where experimental or clinical time is constrained. Both research and clinical applications of TMS would significantly benefit from more efficient motor mapping methods. In this study, we introduced a novel approach that prospectively selects coil configurations based on E-field characteristics. Our findings demonstrate that this method can substantially reduce the number of trials required for motor mapping while maintaining high spatial accuracy.

The primary results of our study demonstrate that the FPS algorithm facilitates more efficient motor mapping compared to random sampling. FPS achieved the same accuracy level (95% overlap with the target map) with approximately half the number of stimulation trials required by random sampling. This reduction in trial count not only minimizes the duration of the experiment, but also reduces the burden of participants, which is particularly critical in clinical applications. Furthermore, FPS exhibited greater robustness across participants, as reflected by narrower confidence intervals in both fitting scores and geodesic distances.

In synthetic data analyses, the FPS algorithm also consistently outperformed random sampling. In all scenarios tested, including muscle representations located at the gyral crown, sulcal wall, and ROI edges, FPS demonstrated superior efficiency and spatial accuracy. The advantage of FPS was especially pronounced for representations located at the sulcal wall and ROI edge. The congruence between the synthetic and experimental findings underscores the reliability and generalizability of the FPS method for TMS-based motor mapping.

Another key finding from our synthetic data analyses is that muscle representations can be recovered with comparable efficacy in both the gyral crown and sulcal wall regions (see Figure 2). This is particularly relevant given the ongoing debate regarding the precise loci of initial TMS activation. While some studies suggest that activation primarily occurs at the gyral crown, others propose the sulcal wall as the initial target [21; 23; 24; 25; 49]. Our simulations revealed that although motor representation recovery is less optimal within deeper cortical folds, our method is unbiased toward either the gyral crown or sulcal wall and can recover representations equally well in both regions. The experimental data revealed that the most probable motor representations for the FDI muscle are located in the gyral crown and lip regions (see Figure 4).

A notable strength of the FPS method is its computational simplicity. The algorithm does not require specialized expertise, operates on standard CPU infrastructure without GPU acceleration, and has a computation time comparable to head modeling processes such as those performed using the SimNIBS headreco pipeline [50]. These features make the FPS accessible for a wide range of research and clinical settings. The method is particularly beneficial for preoperative functional cortical mapping in patients [11; 12; 13; 14; 15; 16; 51; 52; 53; 54] and for exploring higher-order cortical functions in cognitive neuroscience studies involving healthy participants [55]. In these applications, streamlining mapping approaches that reduce participation time while maintaining data quality are essential and may, for example, enable functional mapping of complex cognitive processes such as internal world models [56].

An additional strength of our novel method lies in its ability to minimize operator bias when implementing a truly random randomization strategy. Humans typically struggle to generate true randomness; as a result, random sampling in real experimental conditions often introduces biases, such as a tendency to revisit previously selected sites. However, in our study, this bias was effectively mitigated through the use of algorithmically generated random sampling combined with a neuronavigated robotic arm. These findings suggest that the differences between FPS-based and human-based random sampling might be even more pronounced in real-world experimental settings.

Despite these advantages, the FPS method is not without limitations. It relies on neuronavigated, robotic-arm-guided systems to implement pre-generated coil configurations. Without robotic assistance, experimental validation becomes prohibitively time-consuming. In contrast, random sampling can be conducted with neuronavigation alone, without requiring TMS robot infrastructure or prospective E-field modeling. Furthermore, the FPS method still requires programming expertise in Python or MATLAB, which currently limits its applicability in routine clinical settings. Moreover, FPS introduces an additional computational burden, as it requires calculating the E fields for all candidate coil configurations (***C***). In contrast, random sampling requires calculations only for the coil configurations that were actually utilized. Furthermore, the FPS method currently depends on pre-generated coil configurations, limiting its flexibility to navigate to dynamically generated target locations during motor mapping. Future work should explore closed-loop mapping approaches, where coil configurations are iteratively updated online based on prior stimulation outcomes. The main difficulty of such an approach would be an efficient implementation of the simulation of the E-field that is fast enough to be run during online stimulation to inform the next coil configuration. However, such advancement could further optimize mapping efficiency and adaptability significantly. Additionally, the current study utilized *R*^2^-maps to evaluate the goodness of fit; however, this metric may face challenges in distinguishing between closely situated candidate regions. This limitation arises from its inherent assumption of fitting a sigmoid to a single compartment, presuming that a single anatomically restricted location underlies the causal relationship between E-field strength and response strength. Consequently, it disregards the possibility of multi-modal or combinatorial effects. Incorporating more advanced statistical approaches could improve the specificity and robustness of the mapping process. Regardless of these consideration, our findings highlight the potential of the FPS algorithm to significantly enhance the efficiency and robustness of TMS-based motor mapping. By reducing trial counts, improving consistency, and maintaining high spatial accuracy, FPS represents a promising step forward for both clinical and research applications of TMS.

## 6 Acknowledgment

The study was supported by Federal Ministry of Education and Research, Germany (BMBF, 01GQ2205A). This work is part of BrainLinks-BrainTools which is funded by the Federal Ministry of Economics, Science and Arts of Baden-Württemberg within the sustainability programm for projects of the excellence initiative II. The authors acknowledge the support by the state of Baden-Württemberg through the possibility of using the bwHPC (bwUniCluster 2.0) high performance cluster. The authors thank Petro Julkunen for providing the MATLAB implementation of the adaptive threshold hunting algorithm. The academic English of this paper was improved with the assistance of ChatGPT (version 4.1). The experimental design, content, analyses, and interpretations presented in the paper were entirely the work of the authors and not influenced by ChatGPT.

## 7 Authors contribution

Conceptualization: ZT, DLS, MM and AV; Data curation: DLS and ZT; Formal analysis: DLS with the contribution of ZT and MM; Funding acquisition: AV and JB; Investigation: ZT; participant screening: PCR, JS and TD; Methodology: DLS, ZT, and MM; Project administration: ZT, PCR and AV; Software: DLS, SM and ZT; Computational Framework: DLS with the contribution of MM; Supervision: JB, MM and AV; Resources: AV and JB; Visualization: DLS with the contribution of ZT; Writing - original draft: ZT and DLS with the contribution of all authors. All authors discussed the results and contributed to the final manuscript.

## Appendix A Simulation resolution

### A.1 Angular symmetry

In this study we very using a figure-of-eight coil. Thus, the left and right side of the coil are symmetric and a rotation of the coil by 180^*°*^ should result in a similar induced E-field. With this geometric property, the simulations needed to generate a dataset per participant can be cut to half. Differences between two symmetric positions only arise because of asymmetries in power lines and coil cases. To test whether this is reflected in the data, we calculated the mean relative error of the E-field maps and their 180^*°*^ rotated counterparts for each compartment of the ROI. Figure 6 shows that for all compartments the errors is <10%. Thus, we only took into account rotation angles from 0° to 179° for this study.

**Figure 6:**
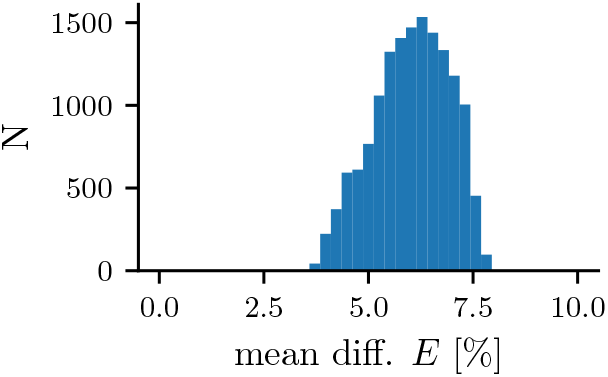
Angular symmetry. Histogram of mean relative difference over 14911 compartments. Differences of 31320 E-fields and their 180° rotated versions are averaged per compartment.

#### A.2 Resolution for FPS

The size of the candidate dataset ***C*** depends heavily on the coil search radius, spatial resolution, and angular resolution of the grid used to generate all possible coil configurations. For example, a search radius of 50mm, angular resolution of 1° between 0 and 179°, and spatial resolution of 1mm results in a size of *N*_*c*_ ≈ 2.800.000 samples. Because E-field simulations are computationally expensive, we examined which resolution is sufficient to have optimal performance of FPS on simulated data.

Figure 7 shows that the goodness of fit is constant for spatial resolutions of 2mm, 3m, 5mm and also for angular resolutions of 10°, 20°, 30°. However, there are differences for various search radii with 30mm yielding the best results. For the spatial and angular resolutions, 5mm and 20^*°*^ were adequate choices. Furthermore, we only considered angles of *a*_*c*_ ∈ [0^*°*^, 180^*°*^) because of the almost symmetric geometry of the coil. This results in a much smaller candidate dataset of *N*_*c*_ = 1017 samples per participant which can be performed in a few hours on a standard computer.

**Figure 7:**
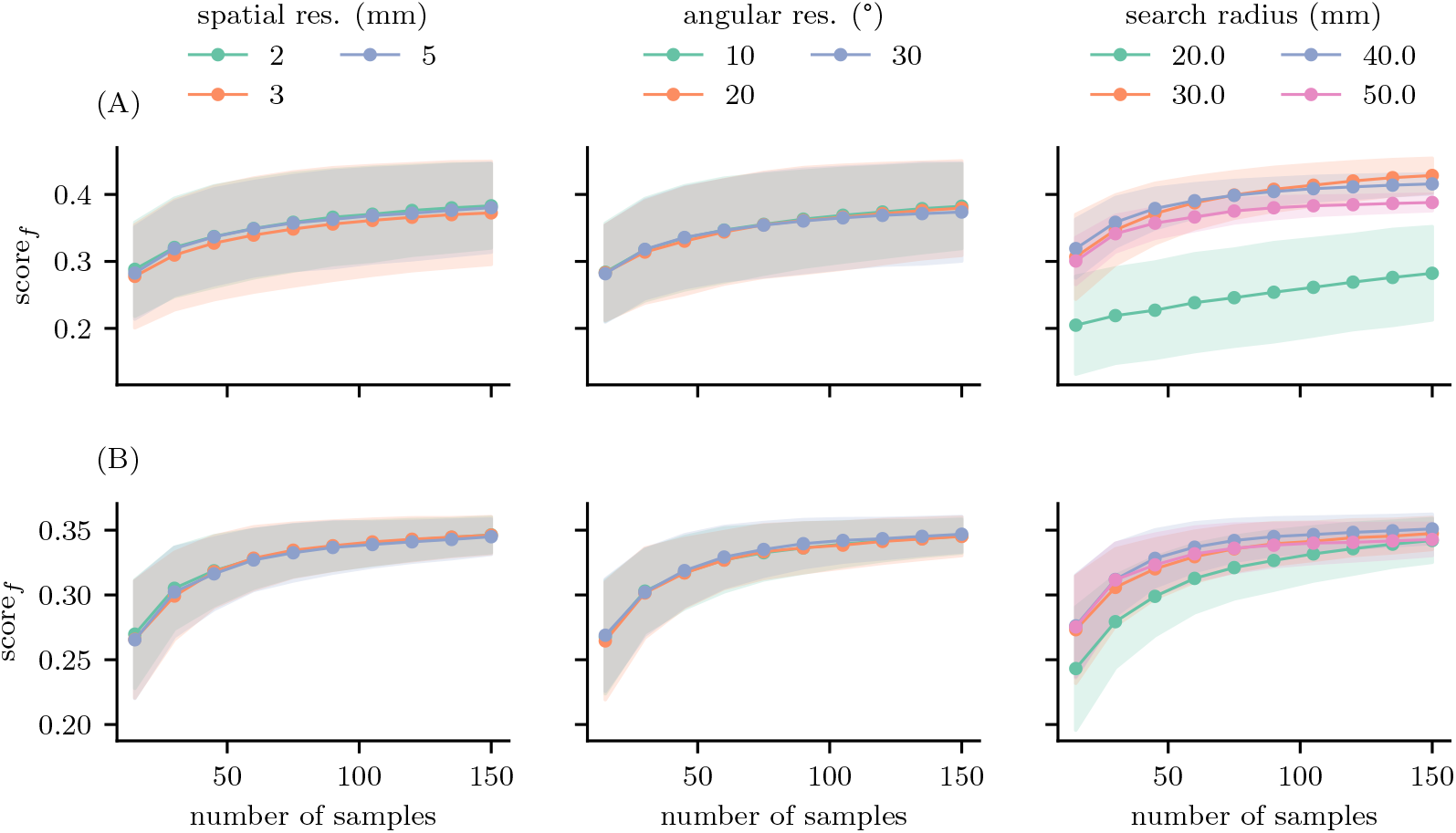
FPS resolution. We varied the spatial resolution, angular resolution, and search radius of the circular grid for generating the candidate E-field dataset for FPS. For each combination of resolution parameters, we tested how well the MGM of the synthetic MEP model could be recovered. We placed the hypothetical muscle representation at (A) sulcal crown in the center of the ROI and (B) a location at the edge of the ROI. Only the search radius has a significant effect on the quality of fit.

## Appendix B Synthetic MEP model parameter sweep

We repeated the experiments conducted in *Section 3*.*1* to study the effect of different MEP model parameters. Figure 8 shows that FPS is outperforming random sampling of the noise amplitude *P*. We also observe, that performance only slowly decreases for larger noise amplitudes, making FPS very robust against MEP amplitude noise.

**Figure 8:**
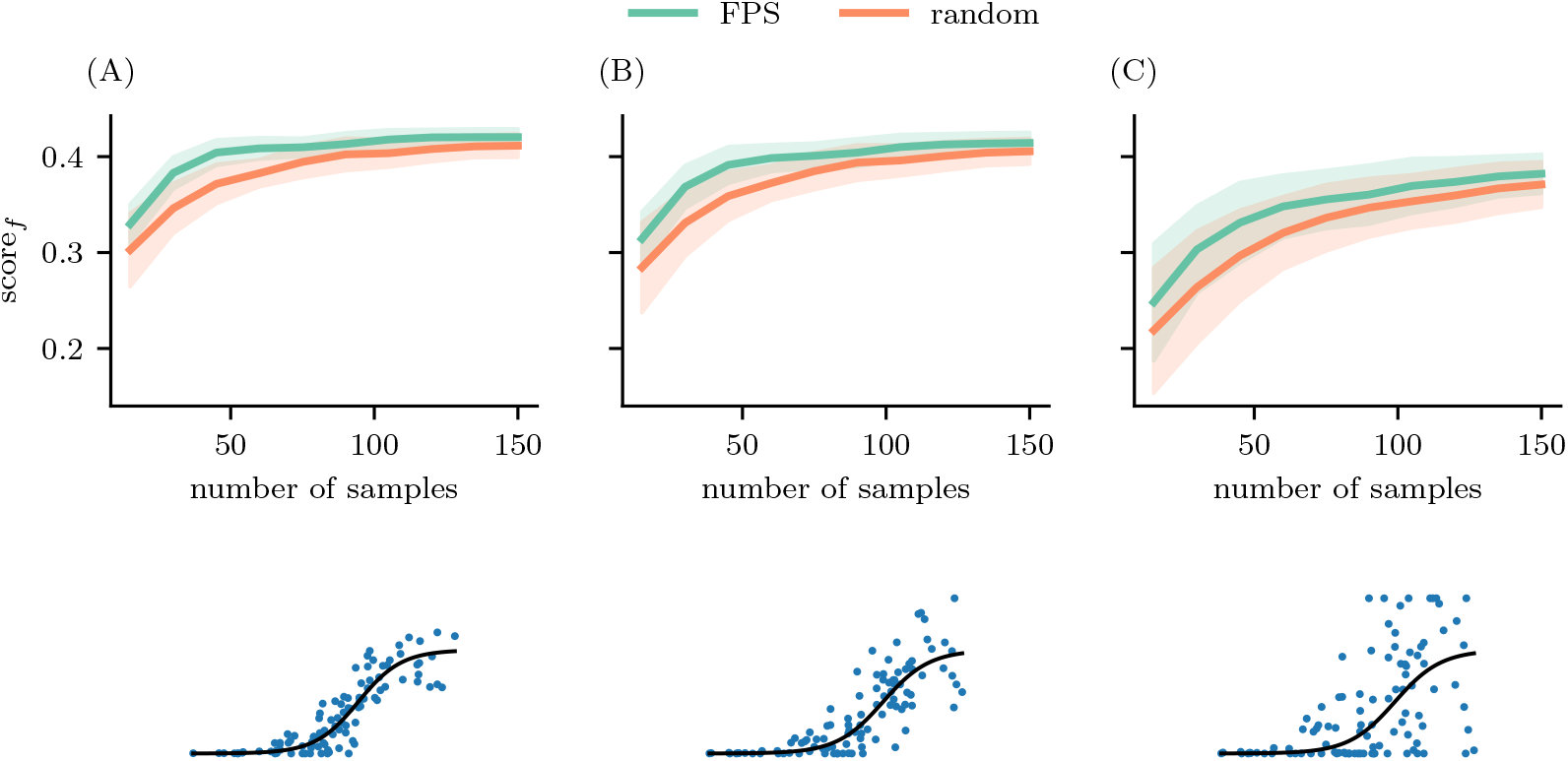
Effect of noise amplitude. We replicated the synthetic data experiment for (A) small, (B) medium, and (C) large noise amplitudes *P* FPS is constantly outperforms random sampling. For larger noise amplitudes the performance of both methods gets worse.

Figure 9 illustrates the effect of muscle representation extension, represented by the MGM standard deviation *σ*. Again, FPS outperforms random sampling for all tested *σ* values. However, the scores indicate that performance drops rapidly with more focalized muscle representations. This hints at the limitation of combining *R*^2^ values into a map as locations close to the true muscle representations are often co-stimulated and therefore also exhibit large *R*^2^-values.

**Figure 9:**
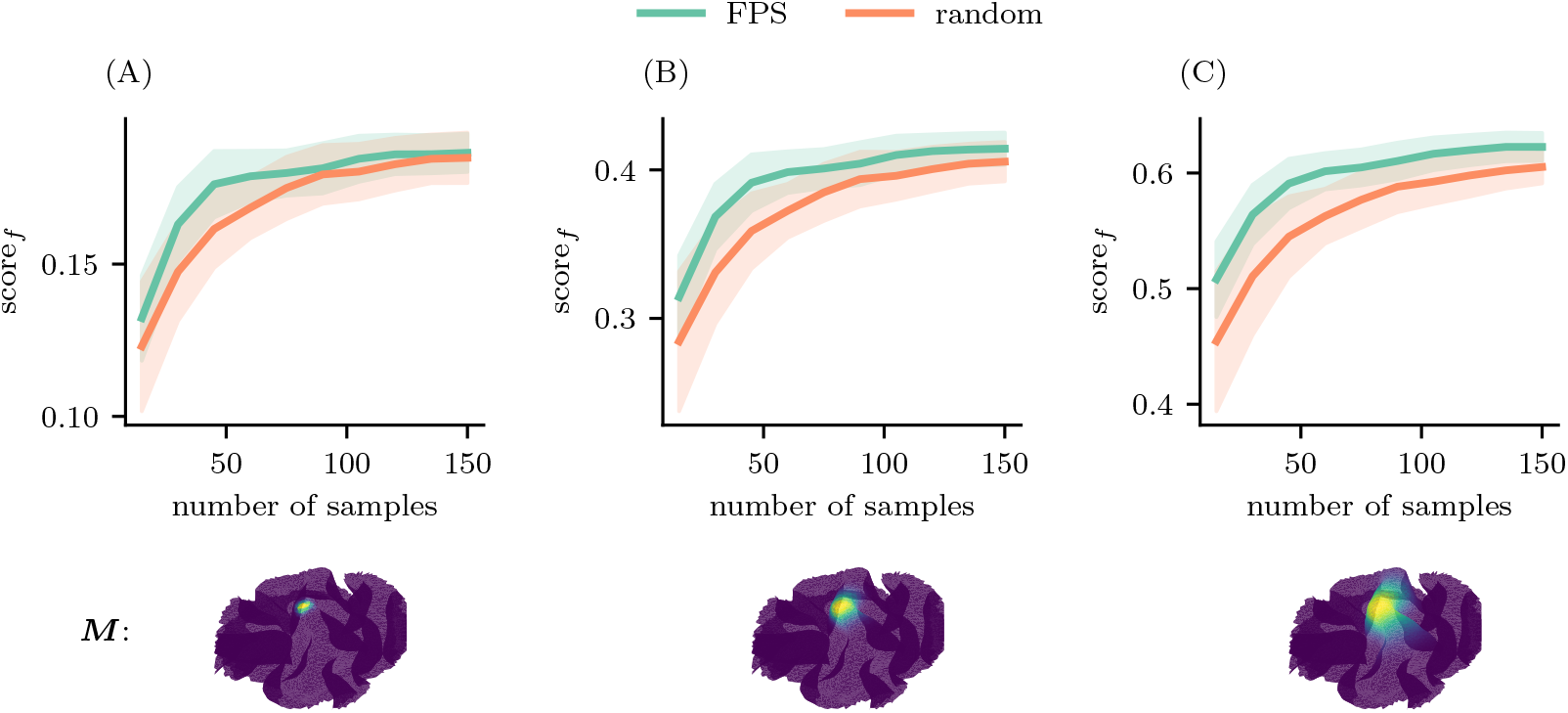
Effect of motor map spread. We replicated the synthetic data experiment for (A) small, (B) medium, and (C) large noise amplitudes *P* FPS is constantly outperforms random sampling. For larger noise amplitudes the performance of both methods gets worse.

Lastly, we tested the effect of the sigmoid lope *k* on the performance as shown in Figure 10. Again, FPS consistently outperforms random sampling. In general, *k* only has neglectable effects on the experiment.

**Figure 10:**
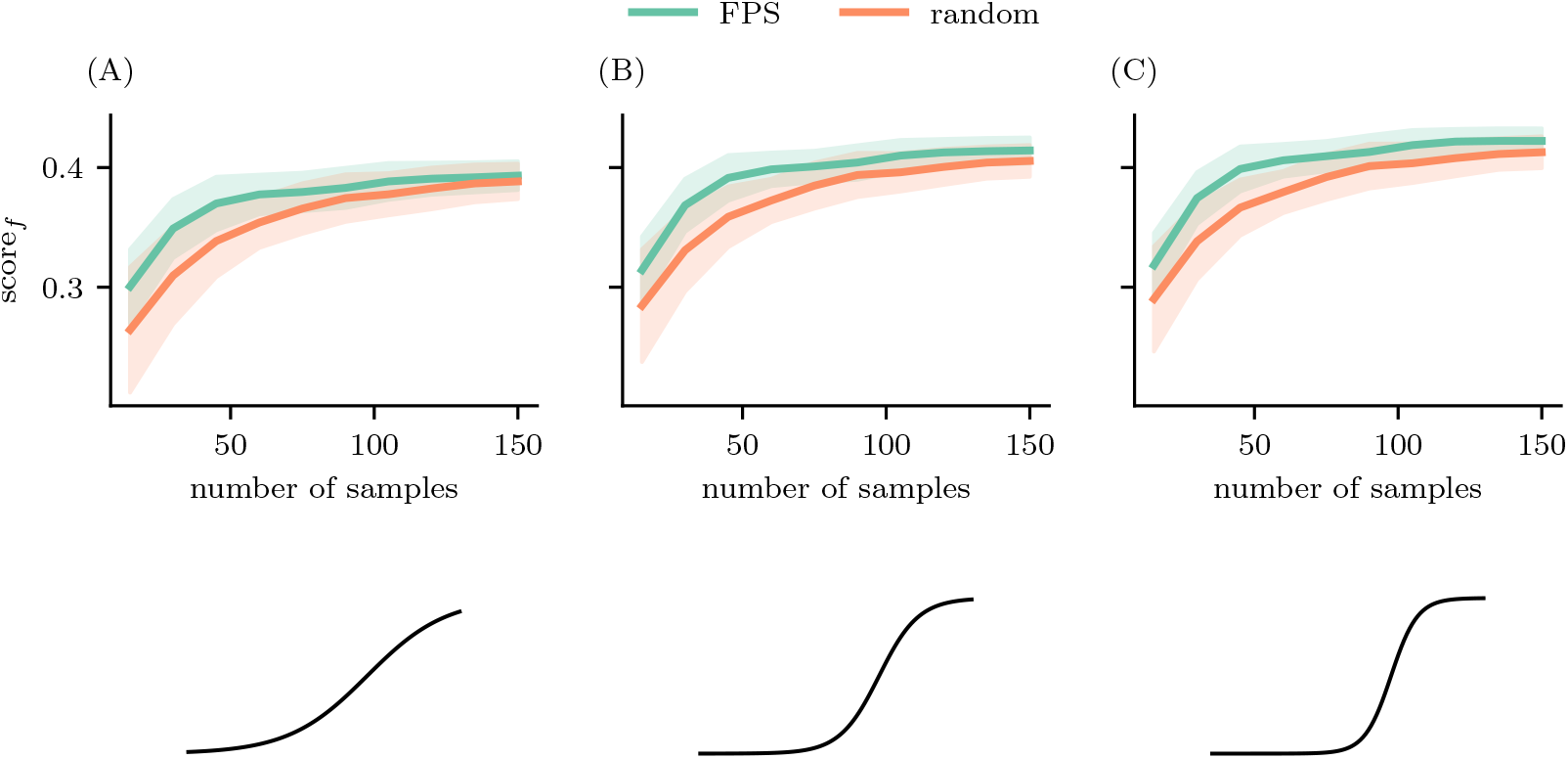
Effect of sigmoid slope. We subsampled a larger dataset from [32] containing MEP Measurements of three muscles for 1005 random coil configurations. For each muscle we measured the fitting score score_*f*_ after every 20 added samples. As ground truth served motor maps fitted with all 1005 samples depicted as insets in the graph. The dots represent the mean across 100 random initializations and the shaded area the standard deviation form the mean. FPS outperforms random sampling for all three muscles, especially in the early sampling phase and has consistently lower variation across initialization

## Appendix C Motor Mapping on previously published data

In their study Nummsen et al. [32] conducted 1005 random stimulations and recorded the MEPs of first dorsal interosseous (FDI), the musculus abductor digiti minimi (ADM), and the musculus abductor pollicis brevis (APB). The data for one participant is publicly available. Thus, there exists a *R*^2^-map for all three muscles retrieved from all 1005 stimulations. We call this the ground truth. We then subsampled the data randomly and with FPS and tested how similar the *R*^2^-maps are for different numbers of subsamples.

Figure 11 shows that pre-selecting E-fields based on FPS yields better results for all three muscles. To reach an overlap of 95%, only 50 FPS samples are needed for all three muscles. With random sampling up to 100 stimulations are needed to reach the same score. This performance gap is closing as the number of samples increases. However, the variance across random initialization (100) of the starting sample leads to larger variance in random sampling than in FPS, making FPS the more robust method. 483.69687pt

**Figure 11:**
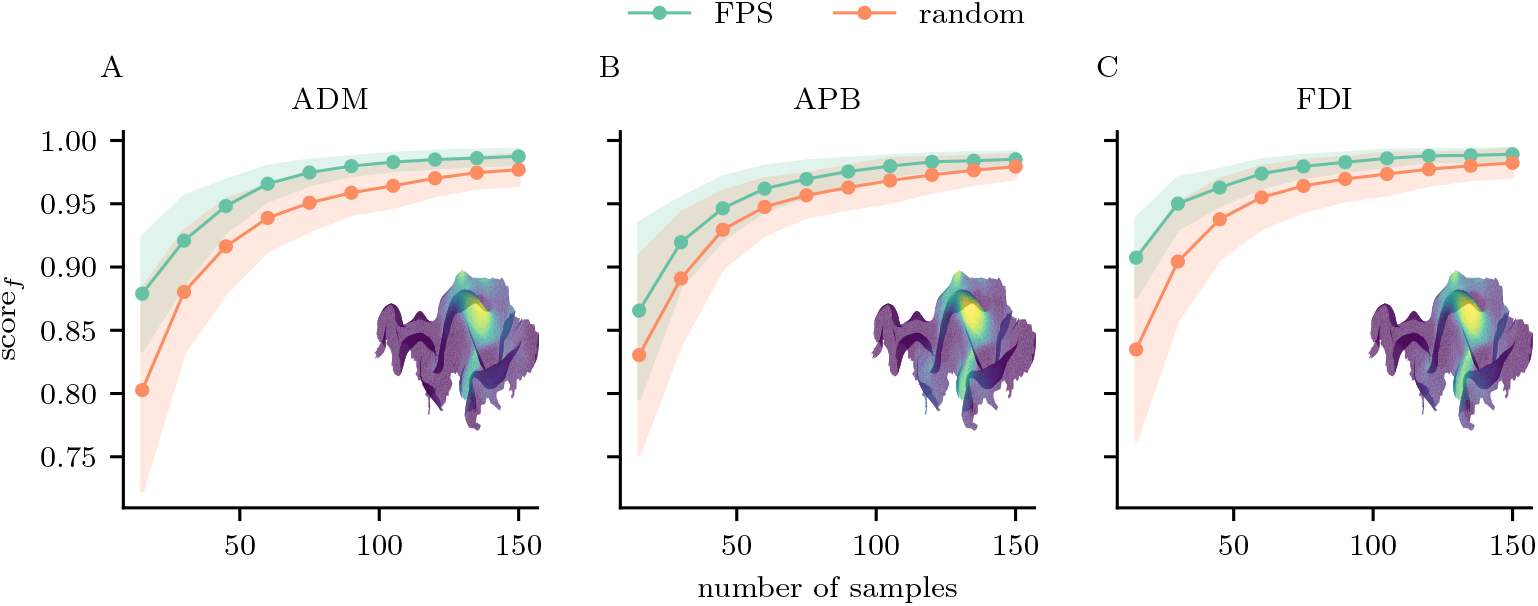
Subsampling larger dataset. We subsampled a larger dataset from [32] containing MEP Measurements of three muscles for 1005 random coil configurations. For each muscle we measured the fitting score score_*f*_ after every 20 added samples. As ground truth served motor maps fitted with all 1005 samples depicted as insets in the graph. The dots represent the mean across 100 random initializations and the shaded area the standard deviation form the mean. FPS outperforms random sampling for all three muscles, especially in the early sampling phase and has consistently lower variation across initialization

**Figure 12:**
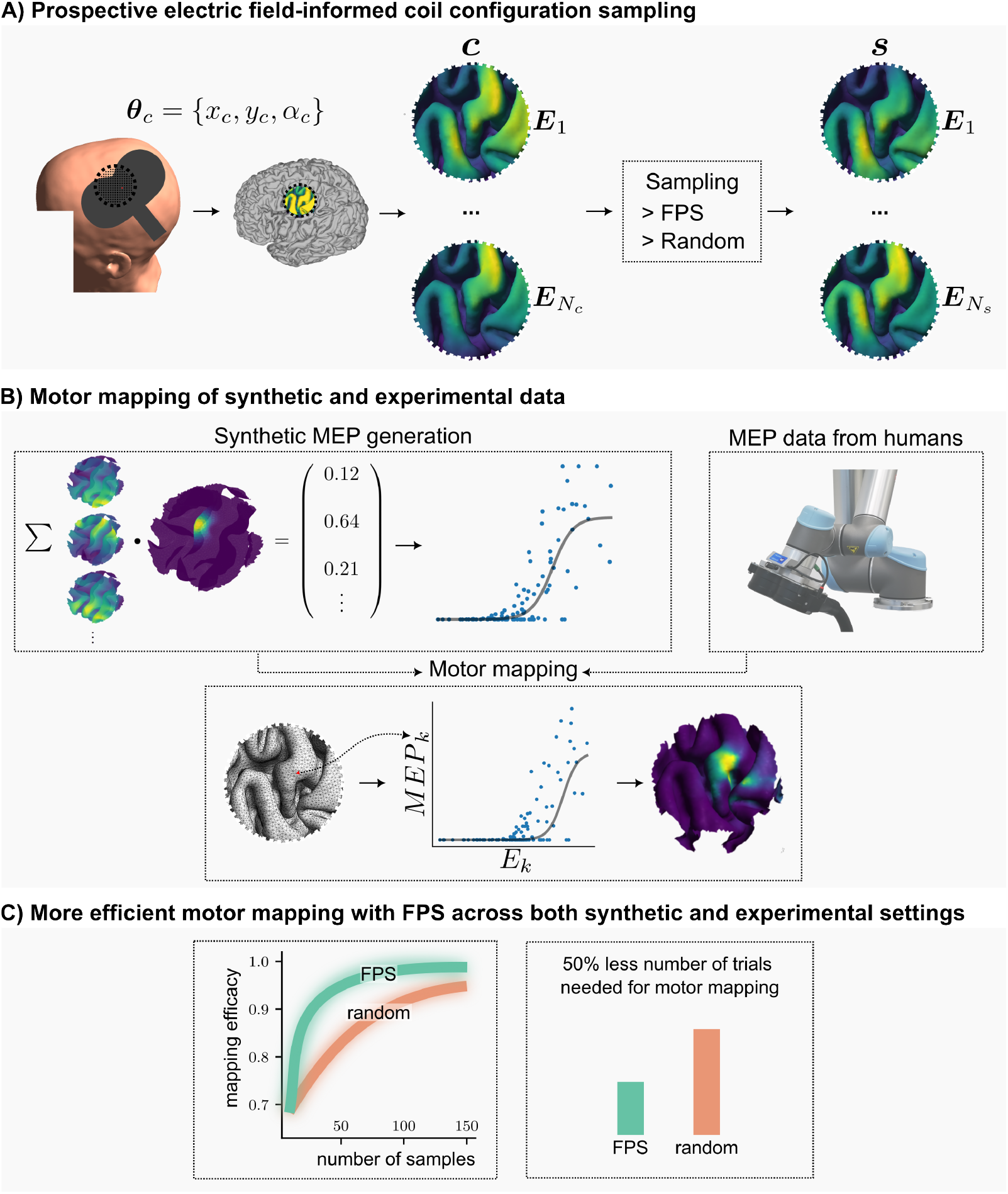
Graphical abstract. A) In this study, we conducted prospective electric field (E-field) simulations for various coil configurations (denoted as ***θ***_*c*_; coil center is denoted as *x*_*c*_ and *y*_*c*_ and coil angle as *α*_*c*_) targeting the motor cortical region of interest (ROI). From the candidate E-field dataset (denoted as ***C***), we selected a subset of coil configurations (denoted as ***S***) using two sampling methods: farthest point sampling (FPS) and random sampling. The FPS method aims to identify coil configurations that are maximally distinct in terms of their E-field characteristics within the ROI. B) The performance of the two sampling algorithms was rigorously evaluated through motor mapping analyses of both synthetic and experimental motor evoked potential (MEP) data. Linear regression was employed to statistically quantify the functional relationship between TMS-induced E-fields within a given mesh compartment (denoted as *E*_*k*_) and trial-specific MEP amplitudes. The resulting motor mapping generates spatial maps that illustrate the goodness of fit between these two variables. C) Data from both synthetic and experimental settings consistently indicate that compared to the random sampling more efficient motor mapping can be achieved with FPS algorithm.

